# Lem2 is essential for cardiac development by maintaining nuclear integrity

**DOI:** 10.1101/2022.02.10.477501

**Authors:** Jacob A. Ross, Nathaly Arcos-Villacis, Edmund Battey, Cornelis Boogerd, Emilie Marhuenda, Didier Hodzic, Fabrice Prin, Tim Mohun, Norman Catibog, Olga Tapia, Larry Gerace, Thomas Iskratsch, Ajay M. Shah, Matthew J. Stroud

**Affiliations:** BHF Centre of Research Excellence, Faculty of Life Sciences & Medicine, King’s College London, London, SE5 9NU, UK; Centre of Human and Applied Physiological Sciences, School of Basic and Medical Biosciences, Faculty of Life Sciences and Medicine, King’s College London, London, SE1 1UL, UK; Hubrecht Institute, Royal Netherlands Academy of Arts and Sciences (KNAW), University Medical Center Utrecht, Utrecht, the Netherlands; Division of Bioengineering, School of Engineering and Materials Science, Queen Mary University of London, London, E1 4NS, UK; Department of Developmental Biology, Washington University School of Medicine, 660S. Euclid Avenue, St Louis, MO, 63110, USA; Crick Advanced Light Microscopy Facility, The Francis Crick Institute, London, NW1 1AT, UK; Department of Molecular Medicine, The Scripps Research Institute, La Jolla, CA 92037, USA; Área de Neurociencias IDIVAL, Facultad de Medicina, Universidad Europea del Atlántico, Santander, Spain

## Abstract

Nuclear envelope integrity is essential for compartmentalisation of nucleus and cytoplasm. Importantly, mutations in nuclear envelope-encoding genes are the second-highest cause of familial dilated cardiomyopathy. One such nuclear envelope protein that causes cardiomyopathy in humans and affects mouse heart development is Lem2. However, its role in mechanically active tissue such as heart remains poorly understood.

We generated mice in which Lem2 was specifically ablated in cardiomyocytes and carried out detailed physiological, tissue and cellular analyses. Importantly, our data showed that Lem2 was essential for cardiac development, and hearts from Lem2 cKO mice were morphologically and transcriptionally underdeveloped. Lem2 cKO hearts displayed high levels of DNA damage, nuclear rupture, and apoptosis. Crucially, we found that these defects were driven by muscle contraction as they were ameliorated by inhibiting myosin contraction and conversely were exacerbated upon myosin activation.

Our data suggest that Lem2 is critical for integrity at the nascent nuclear envelope in fetal hearts, and protects the nucleus from the mechanical forces of muscle contraction. Taken together, these data provide novel insight into mechanisms underlying striated muscle diseases caused by altered nuclear envelope integrity.

## Introduction

The nuclear envelope (NE) in eukaryotic cells separates nuclear and cytoplasmic compartments. Underlying the NE is the nuclear lamina, which consists of Lamins A/C, B1, and B2 (*1*). The nuclear lamina is thought to provide the nucleus with structural rigidity and plays roles in chromatin tethering and gene expression regulation (*2, 3*). The NE is interspersed with Linker of Nucleoskeleton and Cytoskeleton (LINC) complexes that span the double membrane (*4-6*). In striated muscle, such as heart, the LINC complex and its associated proteins have been shown to be essential for normal function (*7-10*). Associated with the LINC complex are the <u>L</u>amina associated peptide 2 (Lap2), <u>E</u>merin, <u>M</u>AN1 (LEM)-domain containing proteins, which also play critical roles in the heart. Indeed, mutations in Emerin, Lap2, and LEM-domain containing 2 (Lem2) are associated with Emery-Dreifuss Muscular Dystrophy with cardiac conduction defects, dilated and arrhythmogenic cardiomyopathies, respectively (*11-15*). LAP2β has been implicated in nuclear envelope repair in a subset of neurons that are under mechanical load during migration (*16*). Therefore we hypothesised that Lem2 might play similar roles to LAP2b in heart that is under constitutive mechanical load.

Lem2 is reported to play roles *in vitro* and in lower organisms. These range from NE repair and resealing after mitosis, maintenance of nuclear shape, regulation of heterochromatin and gene expression, and regulation of mitogen-activated protein kinase (MAPK) signaling (*17-25*). In mice, global loss of Lem2 resulted in embryonic lethality between E10.5-E11.5, with most tissues being substantially smaller in size (*26*). However, the specific roles that Lem2 plays in heart remain poorly understood.

To this end, we aimed to investigate whether Lem2 plays similar roles *in vivo* as those proposed *in vitro* and in lower organisms. Here, we found that Lem2 plays an essential role in heart development. Ablation of Lem2 in cardiomyocytes led to embryonic lethality, with mutant hearts showing marked signs of developmental delay. Transcriptomics revealed enrichment of cell stress pathways and suppression of cardiac developmental pathways, which led us to investigate the cellular effects of Lem2 loss in cardiomyocytes. Lem2-null cardiomyocytes were highly apoptotic, concomitant with both increased DNA damage and micronuclei. Isolation of primary cardiomyocytes revealed aberrant nuclear morphology as well as high levels of nuclear rupture in Lem2-null cells, suggesting that Lem2 plays important roles in maintaining NE integrity. As these cells are highly contractile, we hypothesised that inhibition of mechanical strain on the NE may rescue the nuclear phenotypes. Indeed, we found that inhibiting both myosin contraction and calcium channels attenuated nuclear defects. Conversely, increased activation of sarcomeric myosin exacerbated nuclear damage. Next, we explored roles of Lem2 in adult cardiomyocytes, which we hypothesised would be better adapted to withstand mechanical load, and found no changes in cardiac function or nuclear shape.

Our data suggest that Lem2 is critical for maintaining nuclear integrity in cardiac development in which the NE is in an immature state. Conversely, the presence of a more established nuclear lamina, NE, and LINC complex in adult cardiomyocytes is better able to adapt to both mechanical force and the absence of Lem2.

## Results

### Lem2 is essential for normal heart development

To investigate the role of Lem2 in cardiac function, we generated cardiomyocyte-specific knockout mice using a Cre-LoxP approach by crossing Lem2^f/f^ mice with cardiomyocyte-specific Xenopus laevis myosin light-chain 2 (XMLC2V)-Cre transgenic mice (*27, 28*), hereby designated as Lem2 cKO (SF1A). Lem2 ablation was confirmed with Western blotting and immunofluorescence analyses (SF1B-D). We found that the majority of Lem2 cKO mice did not survive past embryonic day 18.5 (E18.5) (Figure 1A, SF1 E-G), indicating lethality in late fetal stages. Tissue harvesting at E18.5 revealed that the Lem2 cKO hearts were smaller with enlarged atria containing congested blood (Figure 1B), indicating abnormal cardiac function. To investigate this further, we performed High Resolution Episcopic Microscopy (HREM) for 3D organ reconstruction and morphological analyses (*29*). Surprisingly, we did not observe any obvious congenital defects involving abnormal chamber development, which may have explained the fetal demise in Lem2 cKO mice (Figure 1C, D, Supp Figures 2A-C, Supp Movies 1-3). However, ventricle wall thicknesses were slightly reduced at E14.5 (SF2D), which became more pronounced at E16.5 (Figure 1E), suggestive of developmental delay. To test whether vascularisation was affected, a possible indicator of developmental delay, we stained E16.5 hearts with antibodies to smooth muscle 22 alpha (SM22α), but no changes were observed (SF1I).

**Figure 1:**
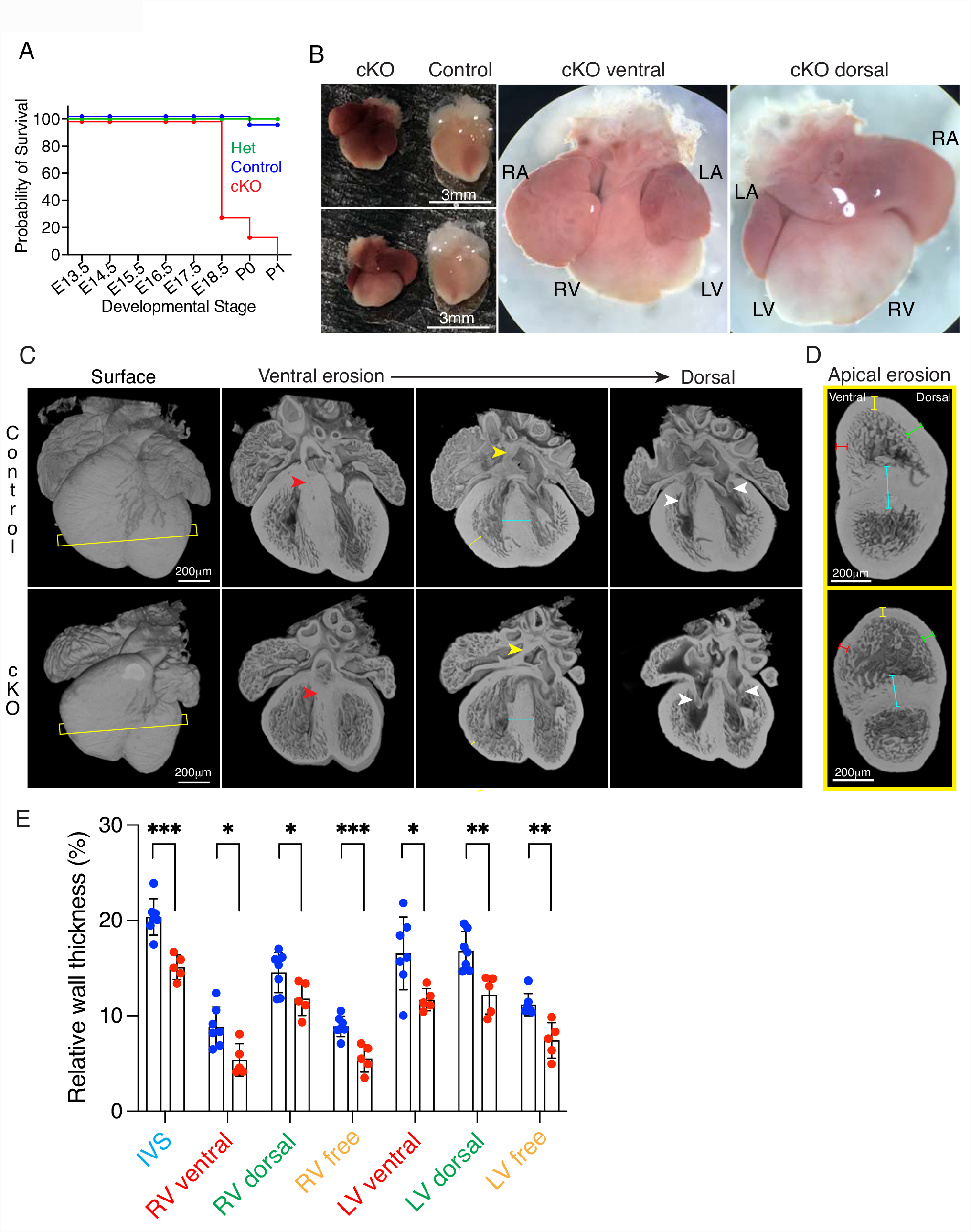
Lem2 is essential for normal heart development. **(A)** Kaplan-Meier survival curve during development from embryonic day 13.5 (E13.5) to postnatal day 1 (P1). Note the fetal demise of Lem2 cardiomyocyte-specific Lem2 knockout (cKO) mice. **(B)** Wholemount images taken prior to birth at E18.5 of cKO and control mice. RA, LA, RV, LV, denote right and left atria, right and left ventricles, respectively. Note the enlarged atria with congested blood in cKO hearts. **(C)** High-resolution episcopic microscopy (HREM) images of whole hearts taken at E16.5 with erosions from the ventral to dorsal surface. Yellow and white arrowheads denote atria septum (AS) and atrio-ventricular (AV) connections, respectively. Yellow slice lines indicate sections imaged in (D). Yellow, blue arrows denote right ventricle freewalls and interventricular septa (IVS), respectively. Note the normal formation of AS and AV connections, but the smaller hearts and underdeveloped wall thicknesses in cKO mice. **(D**,**E)** HREM images eroded from the heart apex showing a mid-ventricle view. Colour-coded lines represent wall thicknesses measured in (E). Blue lines, IVS; Red lines, ventral walls; Green lines, dorsal walls; Yellow lines, freewalls. Note that the walls are all significantly thinner in the cKO compared to controls, indicating delayed cardiac development. *p<0.05, **p<0.01,***p<0.001, two-tailed t-test; n=5-7 mice per genotype. Graphs show mean +/-standard deviation.

### Lem2 ablation activates cell death and inhibits cardiac development pathways

To further assess alterations in cardiac development, we performed RNA-sequencing analysis at E14.5 on control and Lem2 cKO whole hearts. Differential gene expression revealed slight but significant changes to gene expression, with 92 and 49 genes being significantly up- and down-regulated, respectively (Figure 2A). Gene ontology (GO) analysis revealed significant enrichment of the apoptotic cell death pathway and reduction of processes that are key for proper cardiac development, including those involved in morphogenesis, contraction and conduction (Figure 2B, C, F). Upregulated genes included *Gadd45b* and *Gadd45g*, which have been implicated in cell death (*30, 31*), as well as *Fos, Spp1, Arrdc3* (Fig.2D). Downregulated genes included *Rnf207, Kcnq1, Cacng6*, which have been implicated in cardiac conduction and contraction as well as *Adamts6* that regulates heart morphogenesis (*32-35*) (Fig.2G). Importantly, we were able to validate these changes both at mRNA and protein levels (Fig.2E, H). Taken together, these data show that Lem2 is required for cell viability and indispensable for normal cardiac developmental gene expression.

**Figure 2:**
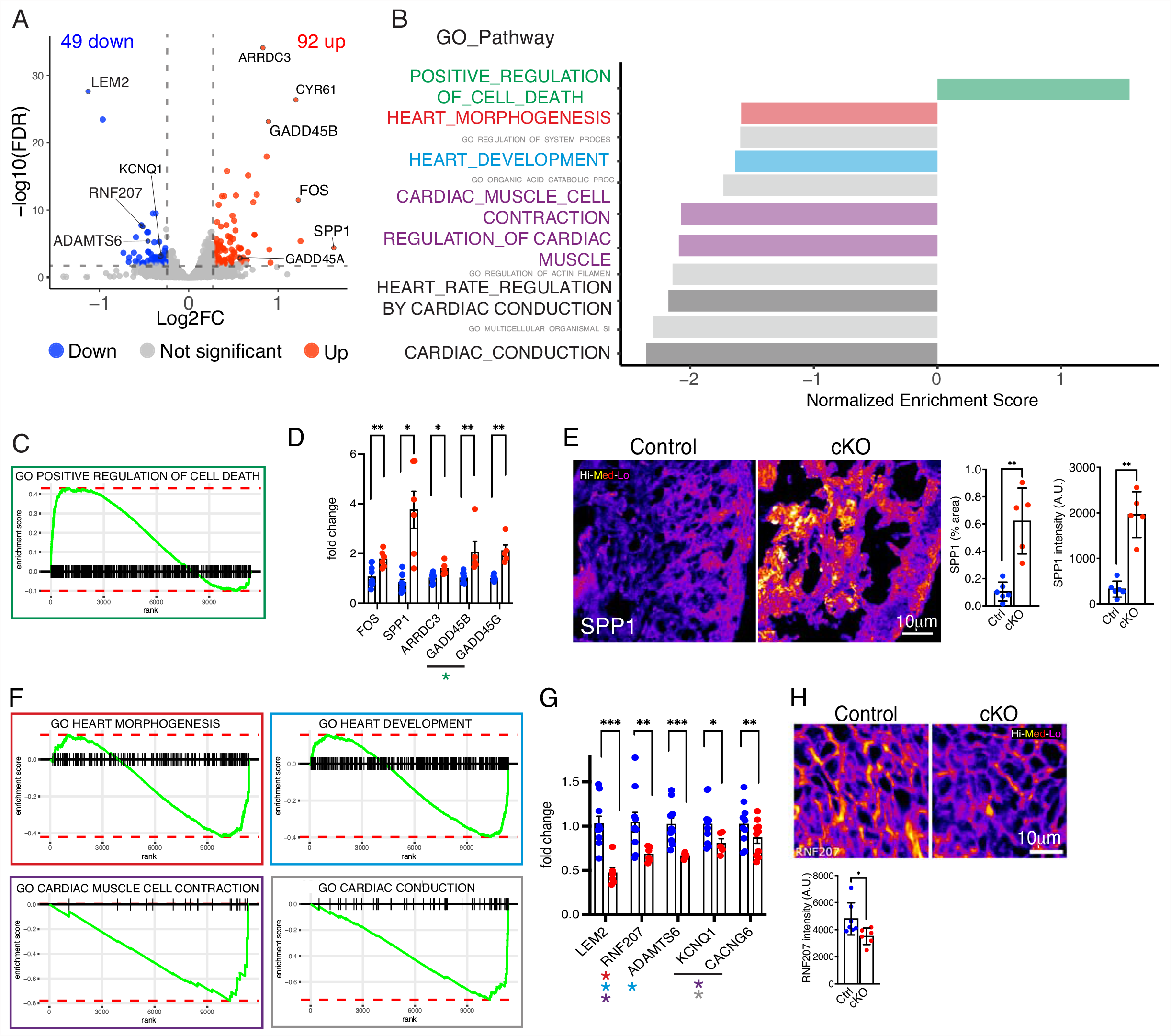
Lem2 ablation activates cell death and inhibits cardiac development pathways. **(A)** Volcano plot showing differentially expressed genes between control and cardiomyocyte-specific Lem2 knockout (cKO) hearts at embryonic day 14.5 (E14.5), from RNA-sequencing data (>1.2 fold change, false discovery rate <0.01). Red and blue dots represent the 92 and 49 genes that were significantly up- and down-regulated, respectively. A selection of qPCR-validated genes are labelled. **(B)** Gene ontology (GO) pathway analysis was performed on differentially expressed genes. Selected terms are highlighted from the top 10 enriched GO-terms. **(C)** GO-term ranking plot shows an over-representation of genes involved in the positive regulation of cell death in cKO hearts. **(D)** RT-qPCR validation of genes identified from RNA-seq as being significantly upregulated in cKO hearts, including cell death genes GADD45A and G. **(E)** Immunofluorescent validation of SPP1, showing increased levels in cKO hearts, further validating the RNA-seq data. **(F)** GO-term ranking plots for shows an under-representation of genes important for cardiac development in cKO hearts. **(G)** RT-qPCR validation of genes identified from RNA-seq as being significantly downregulated in cKO hearts. Note that cardiac genes RNF207, KCNQ1 and CACNG6, as well as Lem2 and ADAMTS6 were significantly decreased. **(H)** Immunofluorescent validation of RNF207, showing decreased levels in cKO hearts, further validating the RNA-seq data. False colour (Fire LUT) used in (E) and (H). *p<0.05, **p<0.01, ***p<0.001, two-tailed t-test; n=4 mice per genotype for RNA-seq; n=7-11 for qPCR validations; n=5 for immunofluorescence validation. Graphs show mean +/-standard deviation.

### Loss of Lem2 results in cardiomyocyte apoptosis, DNA damage accumulation, and micronuclei during cardiac development

To test whether the upregulated cell death pathway resulted in apoptosis, we labelled hearts for cleaved caspase 3, which confirmed high levels of apoptosis in Lem2 cKO hearts between E13.5 and E16.5 (Fig.3A, B, SF3A), consistent with findings in neural tubes of global Lem2 KO mice (*26*). These findings were confirmed using an independent model in which Cre expression was driven by the cardiac troponin T (cTnT) promoter to generate Lem2^f/f;cTnT-Cre/+^ cKO hearts (SF3E, SF1H). Given that Lem2 is a regulator of MAPK signaling, we investigated these pathways, and found that Lem2 cKO hearts displayed elevated ERK1/2, but not p38 activation levels (SF3K). Lem2 has previously been implicated in nuclear envelope resealing and repair, therefore we reasoned that Lem2 ablation may lead to genomic damage and subsequent instability in cardiomyocytes. Indeed, Lem2 cKO hearts showed increased levels of γH2AX (which marks DNA double strand breaks) (*36*) and micronuclei at E14.5 and E16.5 (Figure 3C-F, SF3B-D). Cardiac development is tightly regulated and requires a precise balance between proliferation and apoptosis. We used EdU and anti-phospho-histone H3 staining to mark cells undergoing DNA synthesis and mitosis, respectively. Importantly, proliferation levels were comparable between genotypes (Figure 3G, H, SF3F-I), suggesting that the observed phenotype of smaller hearts and thinner ventricular walls in Lem2 cKO mice is governed by increased apoptosis rather than reduced proliferation.

**Figure 3:**
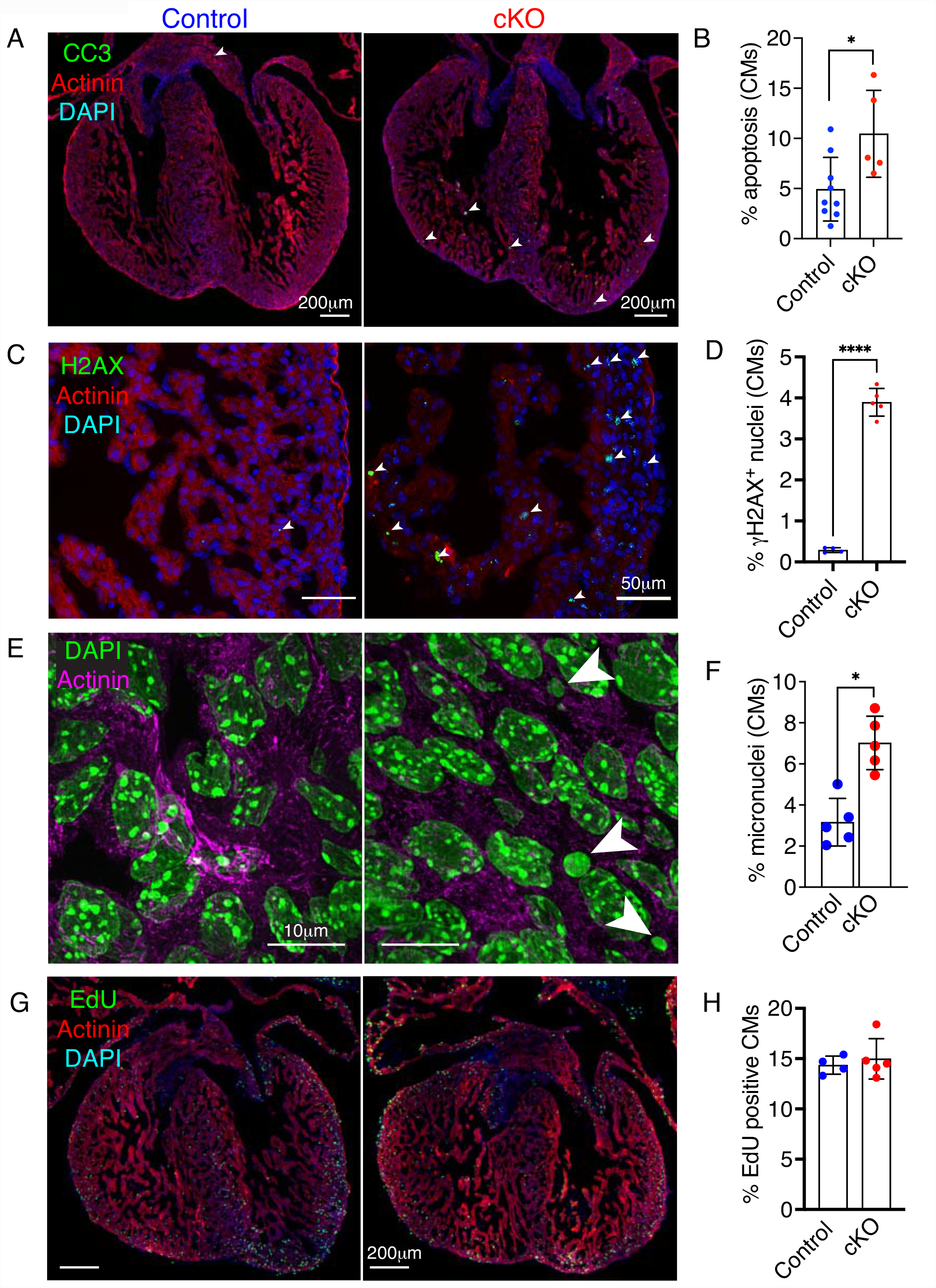
Loss of Lem2 results in cardiomyocyte apoptosis, DNA damage accumulation, and micronuclei during cardiac development. **(A-F)** Fetal hearts at embryonic dayn 14.5 (E14.5) were stained with antibodies raised against cleaved caspase 3 (to mark apoptotic cells) (A); γH2aX (DNA double strand breaks) (C); and sarcomeric α-actinin and DAPI (nuclei). Note the strong increase in cardiomyocytes (CMs) positive for apoptosis,γH2aX, and micronuclei (arrowheads) observed in cardiomyocyte-specific Lem2 knockout (cKO) hearts, quantified in (B, D, F). **(G**,**H)** E14.5 hearts labelled with EdU to delineate cells undergoing DNA synthesis, quantified in cardiomyocytes (H). Note that no changes in EdU incorporation were observed. *p<0.05, ****p<0.0001, two tailed t-test; n=4-8 biological replicates per genotype, with 500-1000 cells quantified/replicate. Graphs show mean +/-standard deviation.

### Cardiomyocyte nuclei lacking Lem2 are aberrantly shaped and more susceptible to mechanical rupture

After observing increased apoptosis, DNA damage, and micronuclei *in vivo*, we next sought to understand the underlying mechanisms using cell biology that is amenable to drug treatments. We plated cardiomyocytes on soft, 6kPa PDMS hydrogels to mimic *in vivo* heart stiffness (*37-39*). Strikingly, we observed aberrant nuclear shape and enlarged nuclear area in Lem2 cKO cardiomyocytes compared to control cells (Figure 4A, B, C). To further assess NE integrity, we transiently expressed catalytically inactive cyclic guanosine monophosphate-adenosine monophosphate synthase (cGAS) fused to an mScarlet fluorescent tag in Lem2 cKO cardiomyocytes to mark cytoplasmic chromatin exposure, hence nuclear rupture events (*40, 41*). Notably, we saw a significantly increased incidence of nuclear rupture in Lem2 cKO cells compared to control, demonstrating an important role for Lem2 in maintaining nuclear integrity (Figure 4D). Importantly, similar observations were made when cells were plated on glass, which is much stiffer than hydrogel (Figure 4B-D), suggesting that the phenotype was independent of forces from the external environment.

**Figure 4:**
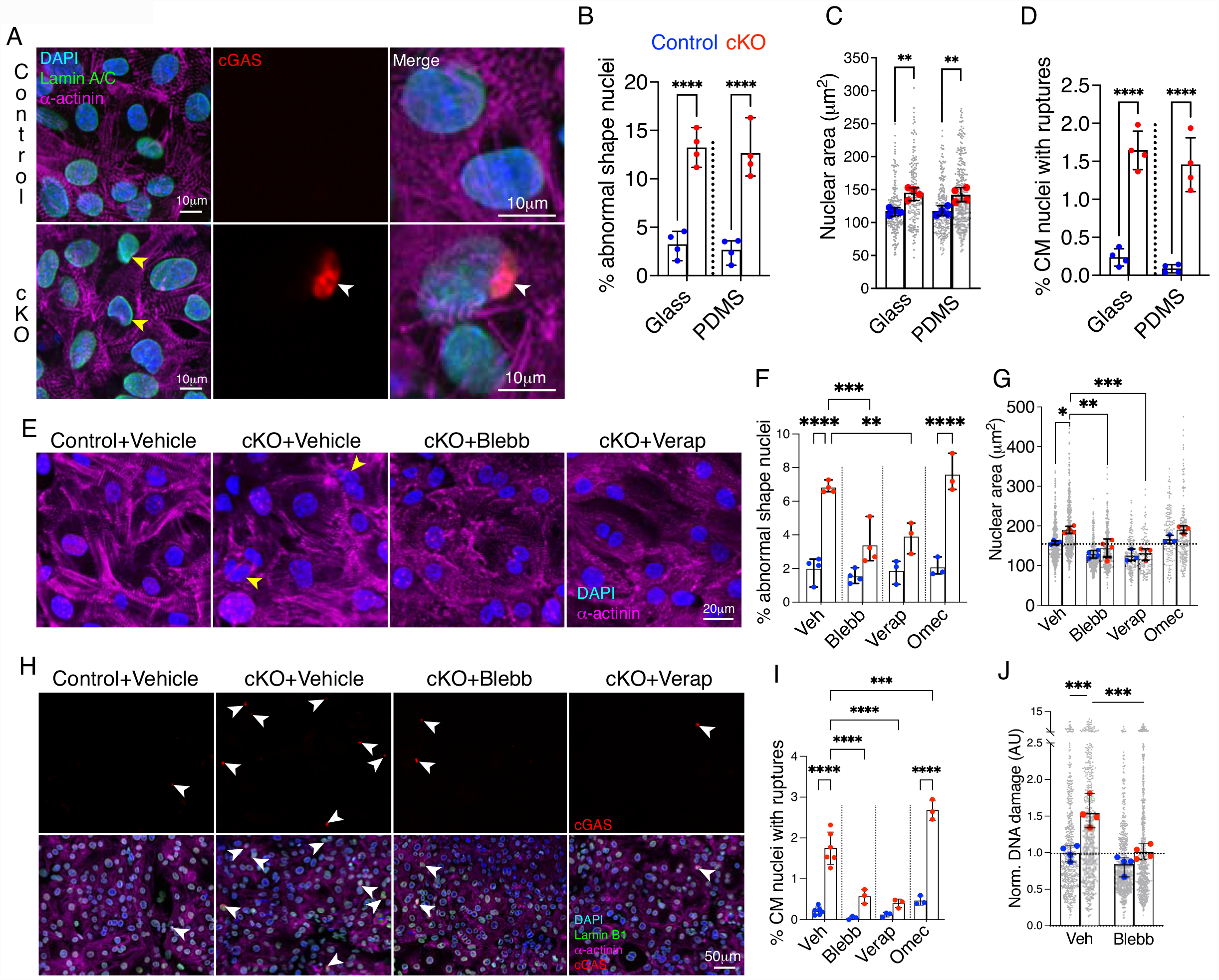
Fetal cardiomyocyte nuclei lacking Lem2 are aberrantly shaped and more susceptible to mechanical rupture and DNA damage. **(A-D)** Fetal cardiomyocytes from control and cardiomyocyte-specific Lem2 knockout (cKO) mice expressing a cGAS-mScarlet probe were isolated and plated on soft, 6kPa PDMS-coated coverslips to mimic stiffnesses encountered in vivo. Cells were stained for DAPI (nuclei), sarcomeric α-actinin (cardiomyocytes) and Lamin B1 (nuclear lamina). Note the aberrant nuclear shapes (yellow arrows in A), larger nuclear sizes and increased nuclear rupture frequency (white arrow in A) observed in the cKO compared to control cardiomyocytes, quantified in (B-D). **(E-I)** Fetal cardiomyocytes from control and cKO mice expressing cGAS probe were plated on glass and treated with blebbistatin (Blebb) or verapamil (Verap) to inhibit muscle contraction or with Omecamtiv mecarbil (Omec) to stimulate contraction. Nuclear shapes and rupture were quantified in (F,G,I). Note the striking attenuation of nuclear shape abnormalities and rupture events observed by inhibiting contraction. In contrast, stimulating muscle contraction increased nuclear rupture. **(J)** cKO cardiomyocytes displayed greater levels of DNA damage compared to control. Note that this was attenuated by inhibiting contraction. Large data points show overall means from each experiment, small points show individual nuclear measurements within each experiment. *p<0.05, **p<0.01, ***p<0.001, ****p<0.0001, two-way ANOVA; n=3-5 experiments with 150-300 cells quantified/experiment. Graphs show mean +/-standard deviation.

Next, we sought to understand the mechanisms underlying nuclear rupture. Given that cardiomyocyte nuclei are under constant mechanical load via the LINC complex, which transmits force from the sarcomeres to the nucleoskeleton (Supplementary movie 4), we hypothesised that inhibiting muscle contraction might attenuate nuclear shape defects and ruptures. To this end, we treated cardiomyocytes with para-nitroblebbistatin which inhibits myosin and arrests muscle contraction (*42*). This agent was able to attenuate nuclear shape defects and rupture (Figure 4E-I), suggesting that contractility is the driver of nuclear pathology in Lem2-null cardiomyocytes. Given that para-nitoblebbistatin is a general inhibitor of the myosin class II family, we hypothesised that these nuclear defects could potentially be driven by non-muscle contractility, i.e. myosin forces originating from the non-sarcomeric cytoskeleton (e.g. cortical actin network and/or actin stress fibres) (*43*). Thus we specifically inhibited muscle contraction via a different mechanism using verapamil, a clinically-used drug which inhibits voltage-gated calcium channels to target excitation-contraction coupling. Similarly, verapamil was also able to attenuate aberrant shapes and ruptures in cardiomyocyte nuclei (Figure 4E-I) (*44*). To further test whether nuclear integrity was affected by muscle contraction, we used a cardiac myosin activator, Omecamtiv mecarbil, to stimulate muscle contraction. In agreement with our hypothesis, we observed increased nuclear rupture in Lem2 cKO nuclei after stimulating contraction. Knowing that muscle contraction induced nuclear rupture, we hypothesised that it may also drive DNA damage, as observed *in vivo* (Fig 3). Indeed, increased DNA damage was apparent in cKO cardiomyocytes relative to controls, that was rescuable by inhibiting muscle contraction. Together, these results show that in cardiomyocytes, Lem2 is essential for maintaining nuclear envelope and genome integrity under mechanical load from muscle contractile forces.

To characterise potential changes in NE composition, we used detailed Western blotting (WB) and immunofluorescence analyses. However, no detectable differences in the majority of NE and lamina proteins were observed between genotypes (SF3J,K). These data suggest that the nuclear defects we observe are driven by lack of Lem2, rather than changes to NE composition.

### Lem2 is dispensable for maintaining cardiac function in adult mice

Having established that Lem2 is essential in fetal cardiomyocytes, we aimed to determine whether it plays a similar role in adults. After establishing expression of Lem2 in adult cardiomyocytes using isolated cells and heart sections (Figure 5A, B), we generated Lem2 inducible conditional knockout (iCKO) mice, using a tamoxifen-inducible cardiomyocyte-specific Cre mouse line Tnnt2MerCreMer/+ (*45*).

**Figure 5:**
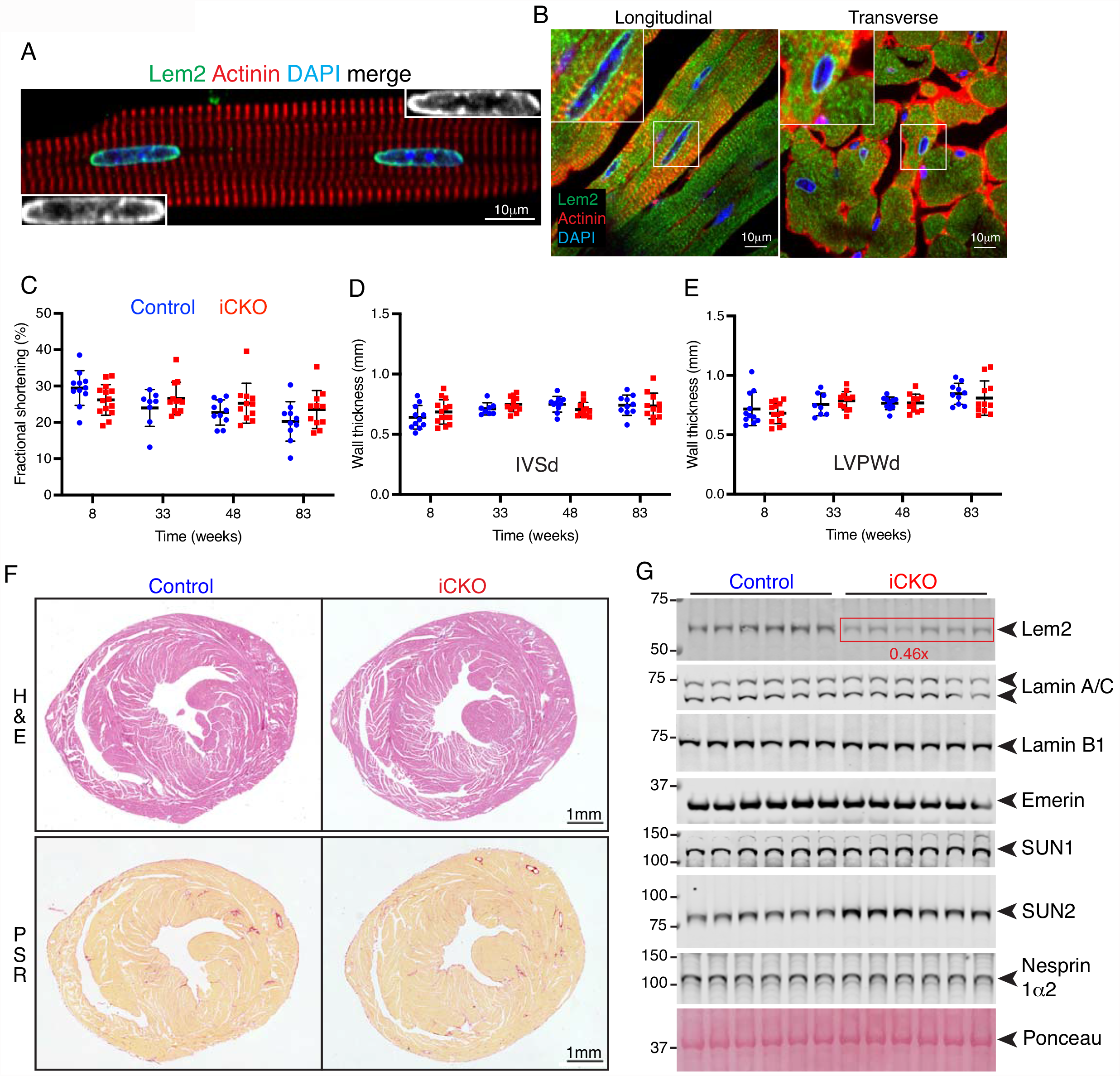
Lem2 is expressed in adult cardiomyocytes, but is dispensable for maintaining cardiac function in adult mice. **(A,B)** Adult cardiomyocytes (A) or longitudinal/transverse sections of adult heart (B) from 8-week-old control mice were stained using antibodies raised against Lem2 (green), sarcomeric α-actinin to denote cardiomyocytes (red) and DNA (blue). Note the discreet localisation of Lem2 to the nuclear envelope in cardiomyocytes. **(C-E)** Echocardiography was performed on Lem2f/f (control) and Lem2f/f;Cre/+ (cardiomyocyte-specific tamoxifen-inducible Lem2 knockout, iCKO) mice at 8 weeks of age, one week prior to tamoxifen injection, and subsequently followed up at 33, 48 and 83 weeks of age. Measured parameters were as follows: (C) Fractional Shortening; (D,E) Inter-Ventricular Septum thickness during diastole (IVSd) and Left Ventricular Posterior Wall thickness during diastole (LVPWd). No differences were observed between control and iCKO. **(F)** Heart sections from 83 week-old mouse hearts stained with haemotoxylin and eosin (H&E) to delineate nuclei and muscle tissue, respectively (top panels); picrosirius red (PSR) to delineate collagen in red (bottom panels). Note there were no differences in morphology or fibrosis between genotypes. **(G)** Western blots were performed on protein extracted from whole hearts from 83 week old mice. Note that as expected, Lem2 levels were reduced in iCKO compared to control, but Lamins A/C, B1, Emerin, SUN1 and Nesprin-1α2 levels were comparable between genotypes. (quantifications shown in Supplementary Figure 4G). n=10-14 mice per genotype for echocardiography, n=6 for Western blot, and n=3 for immunohistochemistry. Graphs show mean +/-standard deviation.

To investigate whether loss of Lem2 specifically affected adult stages of life, we assessed *in vivo* cardiac function in Lem2 iCKO mice. We performed echocardiography at 8 weeks of age, prior to Lem2 ablation at 9 weeks, followed by serial echocardiograms at 33, 48 and 83 weeks of age. Cardiac function and wall thicknesses of Lem2 iCKO mice were comparable to control littermates across all measurements and ages (Fig.5C-E, SF4A-C). Consistently, we observed no changes in expression levels of the fetal gene program and pro-fibrotic markers by quantitative RT-PCR analysis (SF4D), suggestive of no adverse remodelling or fibrosis. Complementary to the echocardiography and gene expression analyses, histological analysis revealed no evidence of morphological defects, changes in extracellular matrix deposition, nor changes in myocyte size (Fig.5F, S4E). Furthermore, there were also no changes in heart weight/tibia length and heart weight/body weight ratios of Lem2 iCKO mice compared to control littermates (S4F). These data suggest that once the adult mouse heart has fully developed, removal of Lem2 does not affect heart function up to 83 weeks of age.

### The majority of NE protein levels are unchanged in Lem2 iCKO hearts and isolated cardiomyocytes

Given the lack of baseline phenotype in the Lem2 iCKO mice, we hypothesised that other nuclear envelope proteins may compensate for Lem2, as shown previously in other systems (*46, 47*). To test this, we performed WBs on whole hearts from control and Lem2 iCKO mice and probed them for a panel of NE and lamina proteins (Fig.5G). As expected, Lem2 levels were reduced to 46% of control levels in Lem2 iCKO (with residual levels of Lem2 likely contributed by non-cardiomyocyte cell types) (SF4G). Other proteins were unchanged, except for the LINC complex protein SUN2, which was increased 1.5-fold compared to control (P<0.01).

Immunofluorescence levels and localisation of most NE proteins were unchanged, with the exception of Lem2 which was absent from the NE in iCKO cardiomyocytes, and a slight increase in SUN2 that is consistent with WBs (Fig.6A-F; S4H,I; S5A, D, E). mRNA transcript levels of *Sun2* (as well as other LINC complex components) were similar between genotypes (SF4J), suggesting that the increase in SUN2 observed in Lem2 iCKO cardiomyocytes is likely regulated at the protein, rather than the gene expression level.

**Figure 6:**
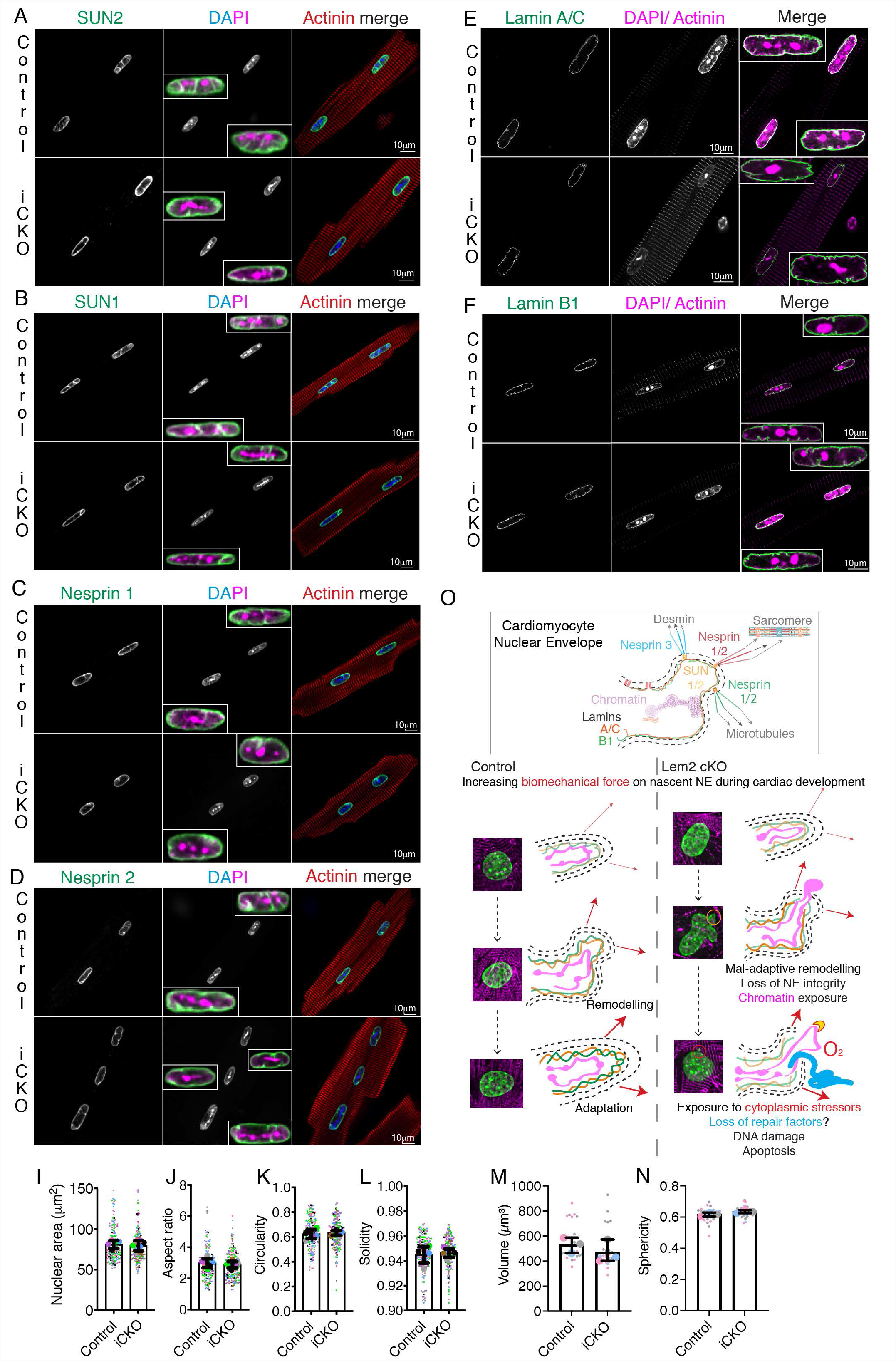
Nuclear shape and nuclear envelope composition are largely unaffected by loss of Lem2 in adult cardiomyocytes. **(A-H)** Adult cardiomyocytes isolated from 8-10-week-old control and cardiomyocyte-specific inducible Lem2 knockout (iCKO) mice were stained using antibodies raised against various nuclear envelope components (green), sarcomeric a-actinin or phalloidin (red) and DNA (blue/ magenta). **(A, B)** SUN2 and 1, respectively, **(C, D)** Nesprin 1 and 2, respectively, **(E, F)** Lamins A/C and B1, respectively, **(G)** Emerin, **(H)** Lem2. **(I-L)** 2D nuclear shape parameters were analysed: nuclear area (I), aspect ratio (J), circularity (K), solidity (L). **(M-N)** 3D shape parameters: nuclear volume (M) and sphericity (N). Conversely, SUN2 levels were increased in Lem2 iCKO isolated adult cardiomyocytes compared to control. Insets show enlarged nuclear envelope compo nents merged with DAPI and maximum intensity projections of Lamin A/C. (**O)** In control cardiomyocytes during fetal heart development, the nascent nuclear envelope (NE) adapts and remodels to mitigate the effects of increases in biomechanical load. Conversely, in Lem2 KO cardiomyocytes, the NE undergoes maladaptive remodelling, which results in nuclear rupture, chromatin exposure and subsequent DNA damage. These ultimately lead to cell death as observed in our cardiomyocyte-specific Lem2 knockout (cKO) model. Conversely, in adult cardiomyocytes that have adapted to withstanding the physical demands on the NE, removal of Lem2 apparently does not affect cardiac function or nuclear shape. Our data suggest that Lem2 is essential for fetal heart develop ment and that removal of Lem2 in adult cardiomyocytes is well tolerated at baseline. (l-o) Large data points represent means from individual mice, small points represent individual nuclear measurements within each mouse. n=3-5 mice/ genotype. Graphs show mean +/-standard deviation.

Knowing that ERK1/2 activation was elevated in fetal Lem2 cKO hearts, we assessed MAPK pathways in adult iCKO cardiomyocytes, but found no differences in activation levels (S5B).

### Nuclear integrity is unaffected in Lem2-null adult cardiomyocytes

Given that we observed alterations to nuclear shape in Lem2 cKO cardiomyocytes, we hypothesised that similar findings might be observed in adulthood. We carried out comprehensive 2D and 3D analyses of cardiomyocyte nuclei, but found no differences in shape parameters between genotypes (Figure 6I-N). Interestingly, we observed invaginations of the nuclear envelope/lamina in both controls and iCKO mice, but to similar extents in both genotypes (Supplementary Movies 5–8). Together these data suggest that Lem2 is not required for normal nuclear morphology in adult mouse cardiomyocytes.

## Discussion

Overall, our data demonstrate that Lem2 is critical for fetal heart development and survival. Lem2 cKO fetal hearts display elevated apoptosis and reduced expression of cardiac developmental genes. Additionally, Lem2-null fetal cardiomyocyte nuclei are highly susceptible to mechanically-induced rupture, resulting in DNA damage, micronuclei and apoptosis. Conversely, ablation of Lem2 in adult cardiomyocytes results in no detectable effects on cardiac function or nuclear shape. Various factors may be able to explain the dichotomy of phenotypes observed between fetal and adult cardiomyocytes including:

Differing Lamin A/C levels between fetal and adult cardiomyocytes: Lamin A/C, which contributes to nuclear rigidity (*48*), is present at low levels in embryogenesis, gradually increasing as cells undergo differentiation and tissues become stiffer (*49*). One could therefore envisage that Lem2 plays a structural role in fetal myocytes prior to the establishment of a more robust, Lamin A/C-containing nuclear lamina. In support of this, Lem2 is expressed in E8.5 mouse embryos, prior to the onset of Lamin A/C expression at E12 (*26, 50, 51*). In contrast to fetal cardiomyocytes, fully differentiated adult cardiomyocytes express higher levels of Lamin A/C, and are therefore better able to mitigate the effect of Lem2 loss. In addition, Lem2 ablation resulted in increased SUN2 levels in adulthood but not in fetal stages, which may be a further compensatory mechanism in fully developed cells.

Differentiation status of fetal compared to adult cardiomyocytes: roles of Lem2 in NE resealing after mitosis point to differing functions in dividing (fetal) versus terminally differentiated (adult) cardiomyocytes (*18, 20, 46*). Our data cannot exclude the possibility that Lem2 plays a role in NE resealing after mitosis, since fetal cardiomyocytes are still dividing. However, we did not observe changes to DNA synthesis or mitosis in Lem2 cKO fetal hearts (Fig.3G, H SF3F-I). Furthermore, the relative subtlety of the cardiac size and morphology defects in fetal Lem2 cKO hearts suggest that Lem2 does not profoundly affect cardiomyocyte division, when compared with other well-established genes involved in this process (*52*). These data favour the hypothesis that the pathophysiological mechanism is likely through defective repair of the NE during interphase.

An intriguing finding of our study was that cardiomyocyte nuclear rupture may be a normal occurrence during cardiac development. In particular, these ruptures are largely the result of forces originating from muscle contraction, which require Lem2-orchestrated repair. We speculate this process may act as a quality control mechanism to ensure nuclear robustness prior to the significant hemodynamic load changes observed postnatally and into adolescence (*36*). Future studies to test this hypothesis would require ablation of Lem2 expression immediately prior to birth to reveal whether these stresses associated with postnatal heart development drive nuclear rupture and if Lem2 is similarly required for this process.

### Working model of Lem2 function in cardiomyocytes

In control cardiomyocytes during fetal heart development, the nascent nuclear envelope adapts and remodels to mitigate the effects of increases in biomechanical load (Figure 6O). Conversely, in Lem2 cKO cardiomyocytes, the nuclear envelope undergoes maladaptive remodelling, which results in gene expression changes, nuclear rupture, chromatin exposure and subsequent DNA damage. These lead to increased cardiomyocyte cell death, which ultimately results in the underdevelopment of the heart as observed in our Lem2 cKO model. Conversely, in adult cardiomyocytes, which harbour nuclei that are stiffer and more resistant to mechanical load, removal of Lem2 apparently does not affect cardiac function or nuclear integrity. Our data suggest that Lem2 is essential for fetal heart development and that removal of Lem2 in adult cardiomyocytes is well tolerated at baseline.

## Acknowledgments

We thank the British Heart Foundation for funding, the Wohl Centre for Cellular Imaging for their support and Camille Charoy for technical assistance.

## Funding

British Heart foundation fellowship FS/17/57/32934 (JR,EB,NA,MJS)

British Heart foundation grant RE/18/2/34213 (JR,EB,NC,AMS,MJS)

British Heart Foundation Chair CH/1999001/11735 (AMS)

National Institutes of Health GM028521 (LG)

## Author contributions

Conceptualization: JR,LG,MJS

Methodology: JR,EB,NA,CB,DH,FB,TM,EM,NC,OT,TI,AMS,MJS

Investigation: JR,EB,NA,CB,FB,TM,NC,MJS

Visualization: JR,NA,MJS

Funding acquisition:MJS

Project administration:MJS

Supervision:MJS

Writing – original draft: JR,MJS

Writing – review & editing: JR,LG,AMS,MJS

## Data and materials availability

All data are available in the main text or the supplementary materials.

## Supplementary Figures

**Supplementary Table 1:** Antibodies

| **Antibody** | **Source** | **Catalogue number** |
| --- | --- | --- |
| Nesprin 1 (WB) | MDA Monoclonal Antibody Resource | MANNES1E Clone 7A12 |
| Nesprin 1 (IF) | Didier Hodzic | n/a |
| Nesprin 2 (IF) | Didier Hodzic | n/a |
| Sun1 (WB) | Abcam | ab103021 |
| Sun1 (IF) | Didier Hodzic | n/a |
| Sun2 (WB) | Epitomics | EPR6557 |
| Sun2 (IF) | Didier Hodzic | n/a |
| Emerin (WB) | Santa Cruz | SC15378 |
| Emerin (IF) | Nova-castra | NCL-Emerin |
| Lamin A/C | Larry Gerace | n/a |
| Lamin B1 | Larry Gerace | n/a |
| Lem2 | Sigma | HPA017340 |
| BAF1 | Abcam | ab129184 |
| SPP1 | Proteintech | 22952-1-AP |
| SM22a | Abcam | ab14106 |
| gamma-H2AX | Cell signalling | 9718S |
| Phospho-histone 3 | Cell signalling | 3377 |
| Cleaved caspase 3 | Cell signalling | 9661 |
| Alpha actinin | Sigma | A7811 |
| RNF207 | Abnova | H00388591-MO2 |
| GAPDH | Santa Cruz | SC32233 |
| Histone 3 | Cell signalling | 4499T |

**Supplementary Table 2:**
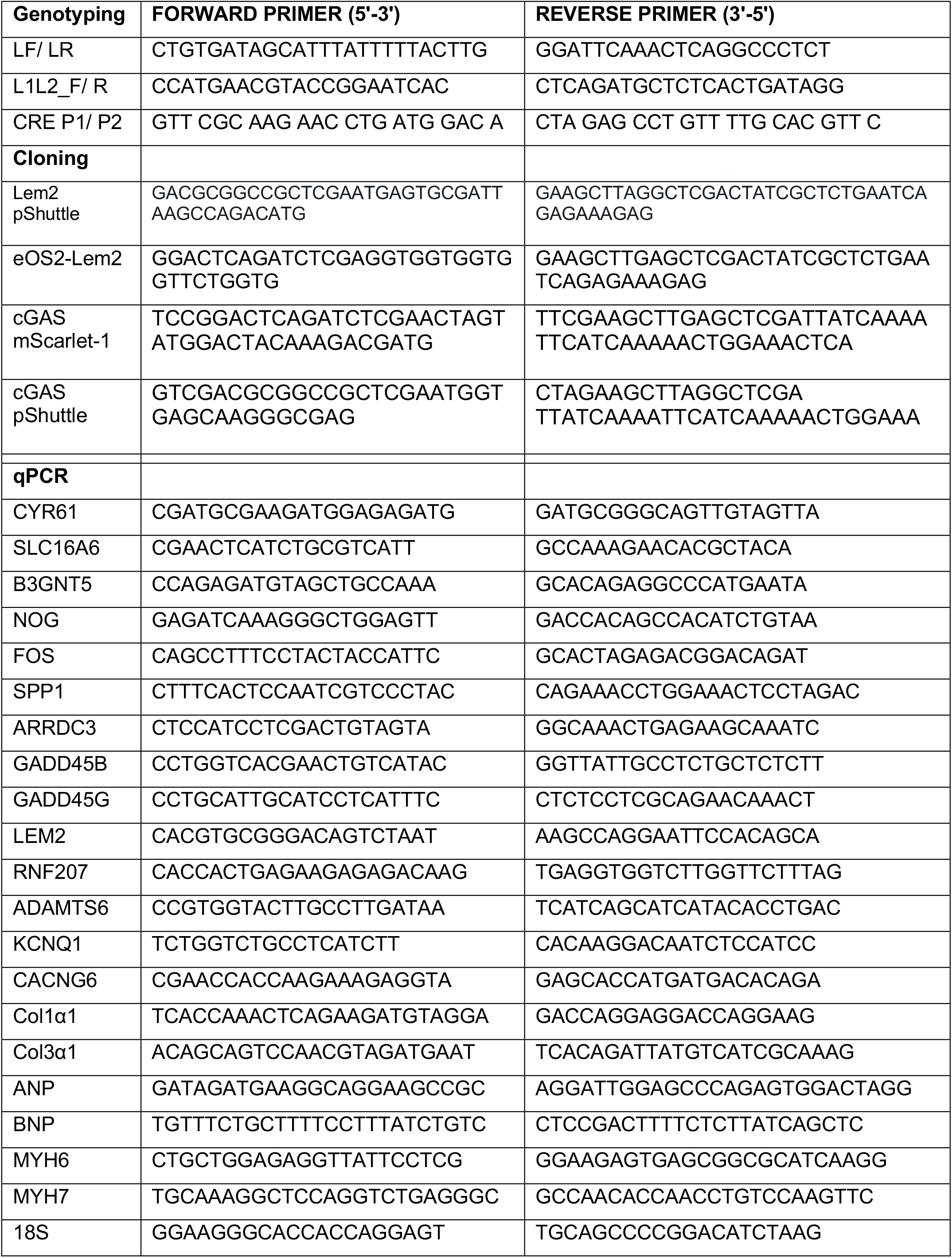
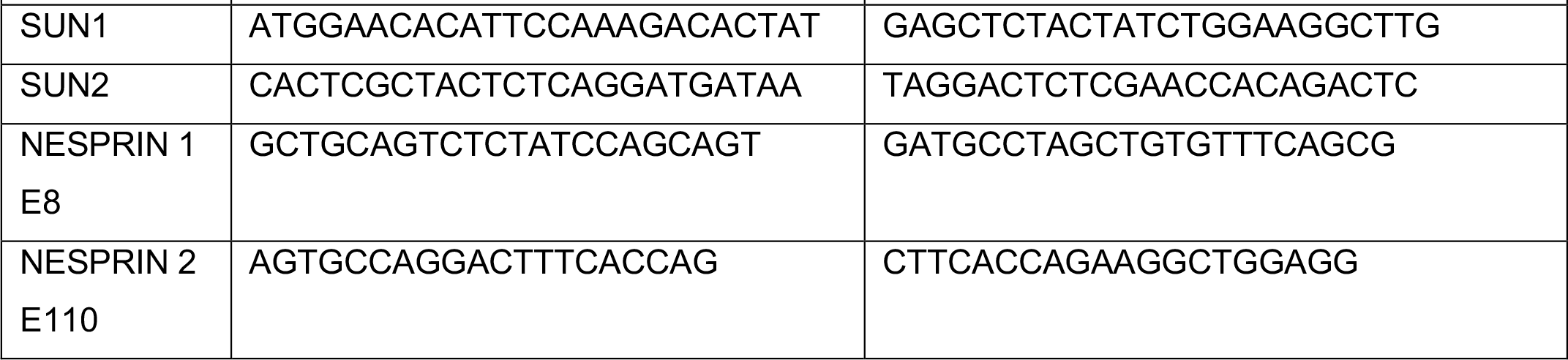
Primers

**Supplementary Figure 1:**
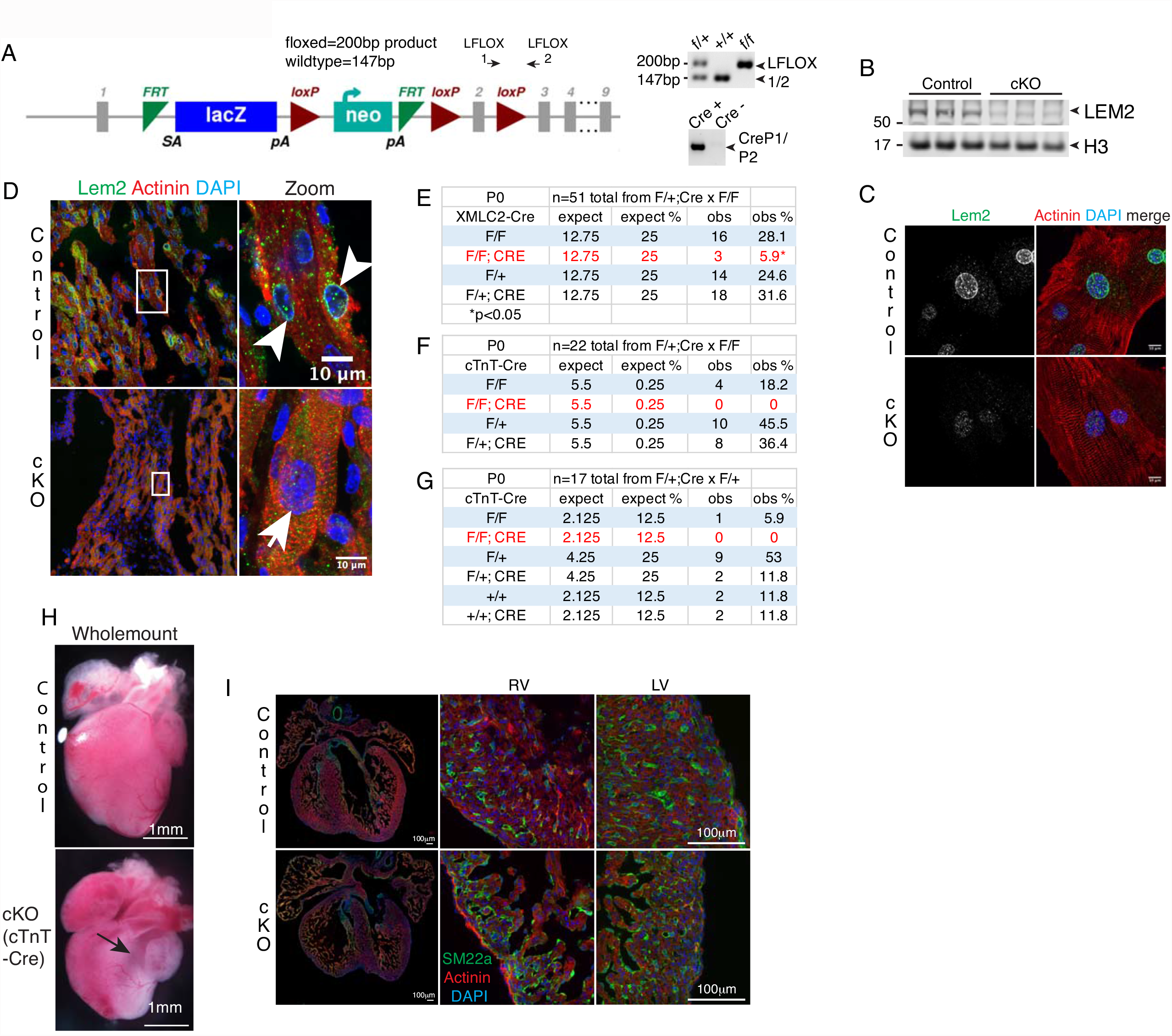
Generation and validation of Lem2 cKO mice. **(A)** Left panel, construct used to generate cardiomyocyte-specific Lem2 knockout (cKO) mice. Arrows above construct indicate primers used for genotyping. Right panel, representative genotyping. **(B)** Western blots from whole hearts at embryonic day 16.5 (E16.5) using an antibody raised against Lem2. Note the reduced level of Lem2 in cKO hearts. **(C**,**D)** Immunofluorescence of isolated cardiomyocytes (C) and heart tissue (D) from fetal hearts stained with antibodies raised against Lem2 and sarcomeric α-actinin (to mark cardiomyocytes). Note the lack of Lem2 staining at the nuclear envelope in cKO cardiomyocytes. **(E-G)** Mendelian ratios at postnatal day 0 from homozygous floxed (Lem2f/f) (E, F) or heterozygous floxed (Lem2f/+) (G) mice crossed with Lem2f/+;XMLC2-Cre/+ or Lem2f/+;cTnT-Cre/+, respectively. Note that mice are not present at the predicted Mendelian ratios of 1 in 4 for (E,F) and 1 in 8 for (G). **(H)** Postnatal day 0 hearts isolated from control and Lem2f/f; cTnt-Cre/+ were whole-mounted. Note the smaller size, almost translucent ventricle wall (black arrow). **(I)** Immunofluorescence on E16.5 heart sections from control and cKO mice using antibodies against SM22a (to denote vasculature) and sarcomeric α-actinin. Note the comparable levels of vascularisation between control and cKO genotypes. RV, LV, right and left ventricles, respectively.

**Supplementary Figure 2:**
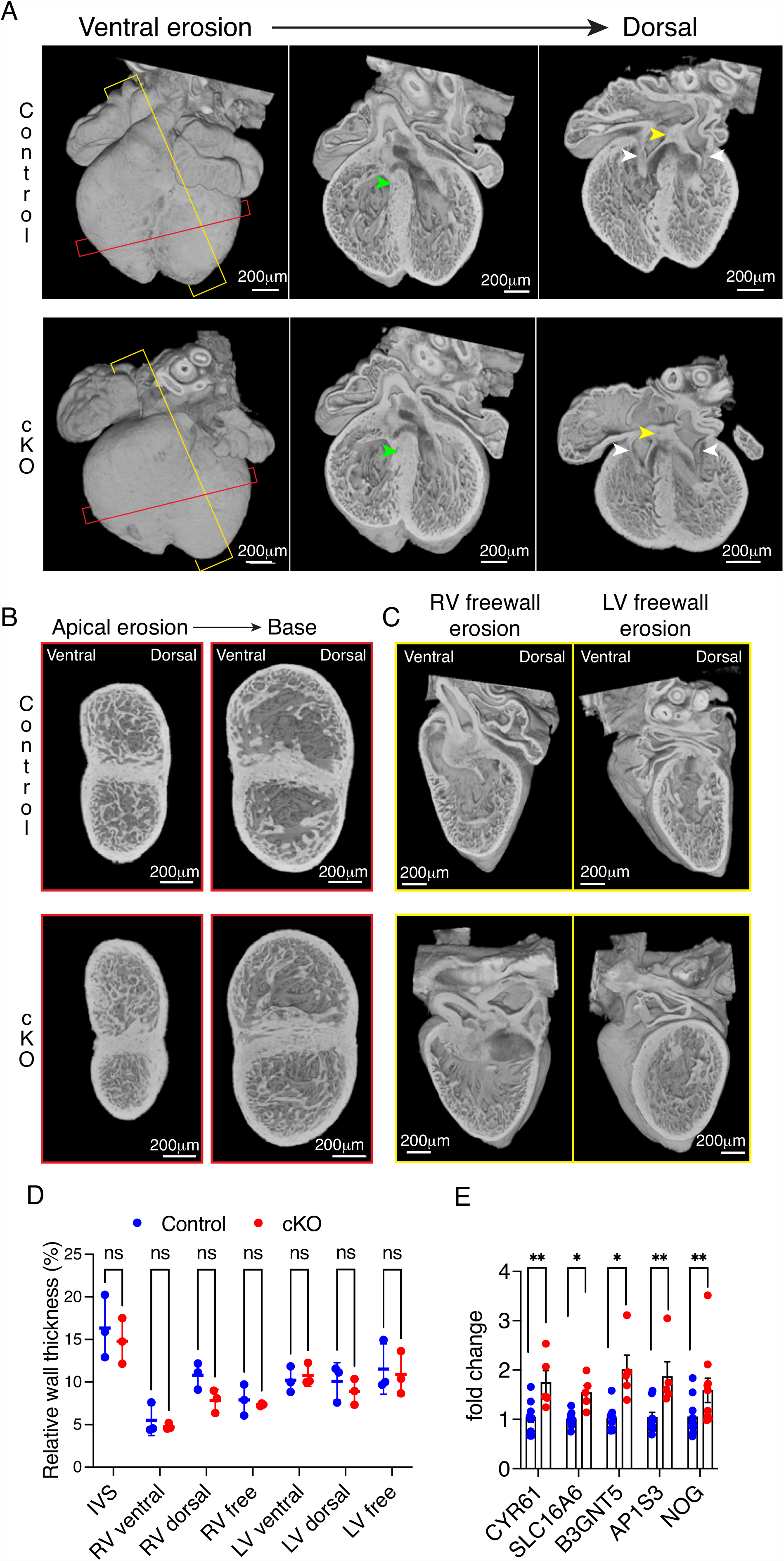
HREM at E14.5. **(A)** High-resolution episcopic microscopy (HREM) images of whole hearts taken at E14.5 with erosions taken from the ventral to dorsal surface. Green, yellow, and white arrowheads denote ventricular septum (VS), atria septum (AS) and atrio-ventricular (AV) connections, respectively. Yellow and red slice lines indicate sections imaged in (B) and (C), respectively. Note the normal formation of VS, AS, and AV connections. **(B**,**D)** HREM images eroded from the heart apex showing mid-ventricle and apical views, quantified in (D). **(C)** HREM images eroded from the right and left freewalls showing a mid-ventricle view. Note that the walls are slightly thinner in the cardiomyocyte-specific Lem2 knockout (cKO) hearts compared to controls, indicating a slight delay of cardiac development. **(E)** RT-qPCR validation of genes identified as significantly upregulated from RNA-sequencing data in Figure 2. *p<0.05, **p<0.01, two tailed t-test; n=4-5 mice per genotype. Graphs show mean +/-standard deviation.

**Supplementary Figure 3:**
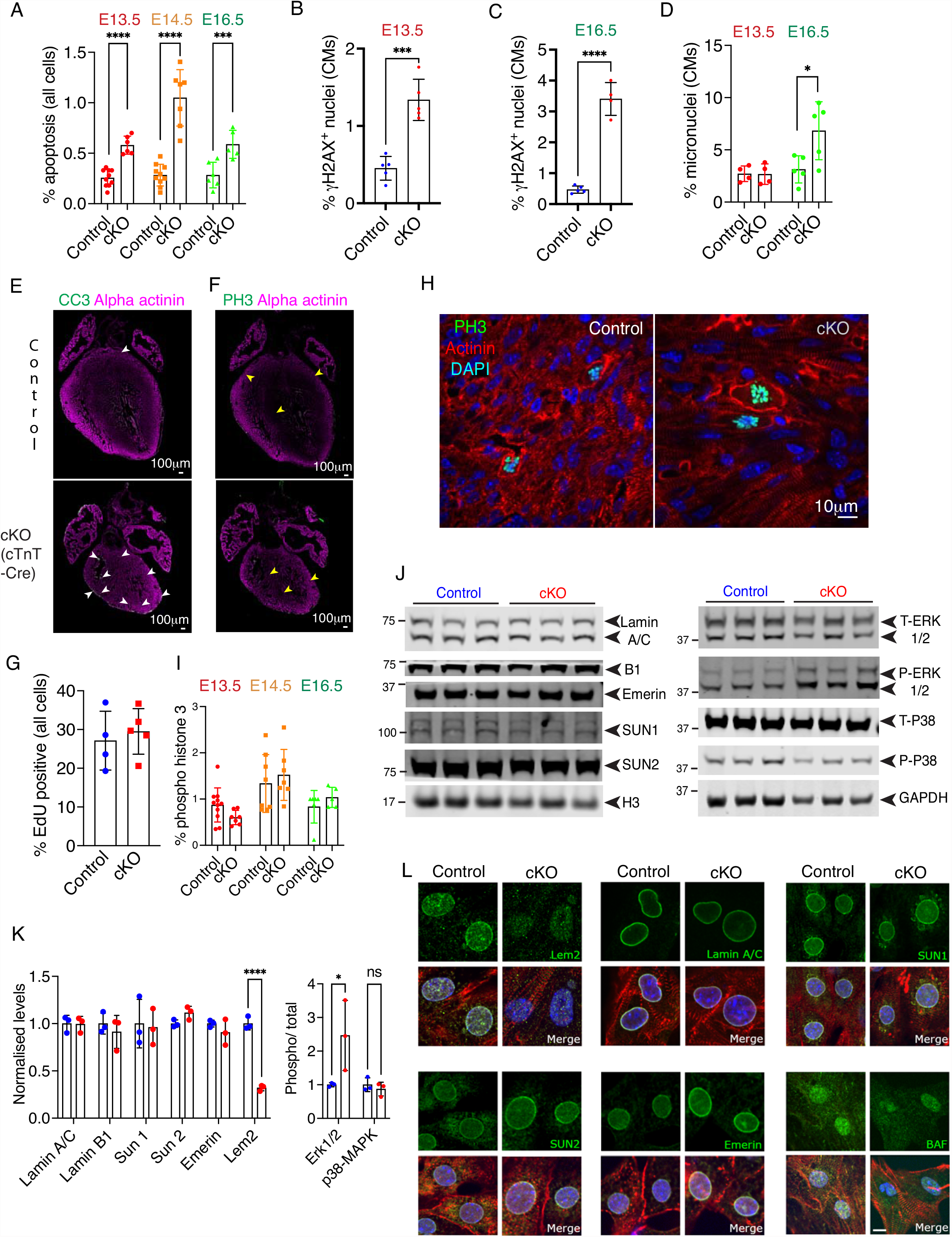
Phenotyping of proliferation, cell death, DNA damage, and micronuclei at different stages of cardiac development. **(A)** Embryonic day 13.5-16.5 (E13.5-16.5) hearts were stained against cleaved caspase 3 to denote apoptotic cells and quantified. Note the increase in apoptosis throughout development in cardiomyocyte-specific Lem2 knockout (cKO) mice compared to control. **(B**,**C)** E13.5 and E16.5 hearts were stained for γH2aX to mark DNA damage in cardiomyocytes. Note the significantly increased levels of DNA damage. **(D)** Micronuclei were quantified in E13.5 and E16.5 hearts. Note the significant increase at E16.5. **(E**,**F)** Postnatal day 0 hearts isolated from control and Lem2f/f; cTnt-Cre/+ were sectioned and stained against cleaved caspase 3 (CC3) to mark apoptosis (white arrowheads) and phospho-histone 3 (PH3) to mark mitosis (yellow arrowheads). Note the increased apoptosis, but comparable levels of mitosis in cKO hearts, which phenocopies the earlier results observed with the Lem2f/f; XMLC2V-Cre/+ mice. **(G)** E14.5 hearts were labelled with EdU to denote cells undergoing DNA synthesis. **(H**,**I)** E13.5-16.5 hearts were stained against phospho-histone 3 (PH3) to denote mitotic cells and quantified in (I). Note that no changes in mitosis were observed throughout development. **(J**,**K)** Western blots from E16.5 whole hearts stained with various nuclear envelope (NE) proteins, quantified in (K). **(L)** Immunofluorescence of isolated cardiomyocytes at E16.5 were stained for various NE components. Note the reduced signal of Lem2 in cKO cardiomyocytes. *p<0.05, ***p<0.001, ****p<0.0001, two tailed t-test; n=4-8 biological replicates per genotype, with 500-1000 cells quantified/replicate for immunohistochemistry; n=3 hearts for Western blot. Graphs show mean +/-standard deviation.

**Supplementary Figure 4:**
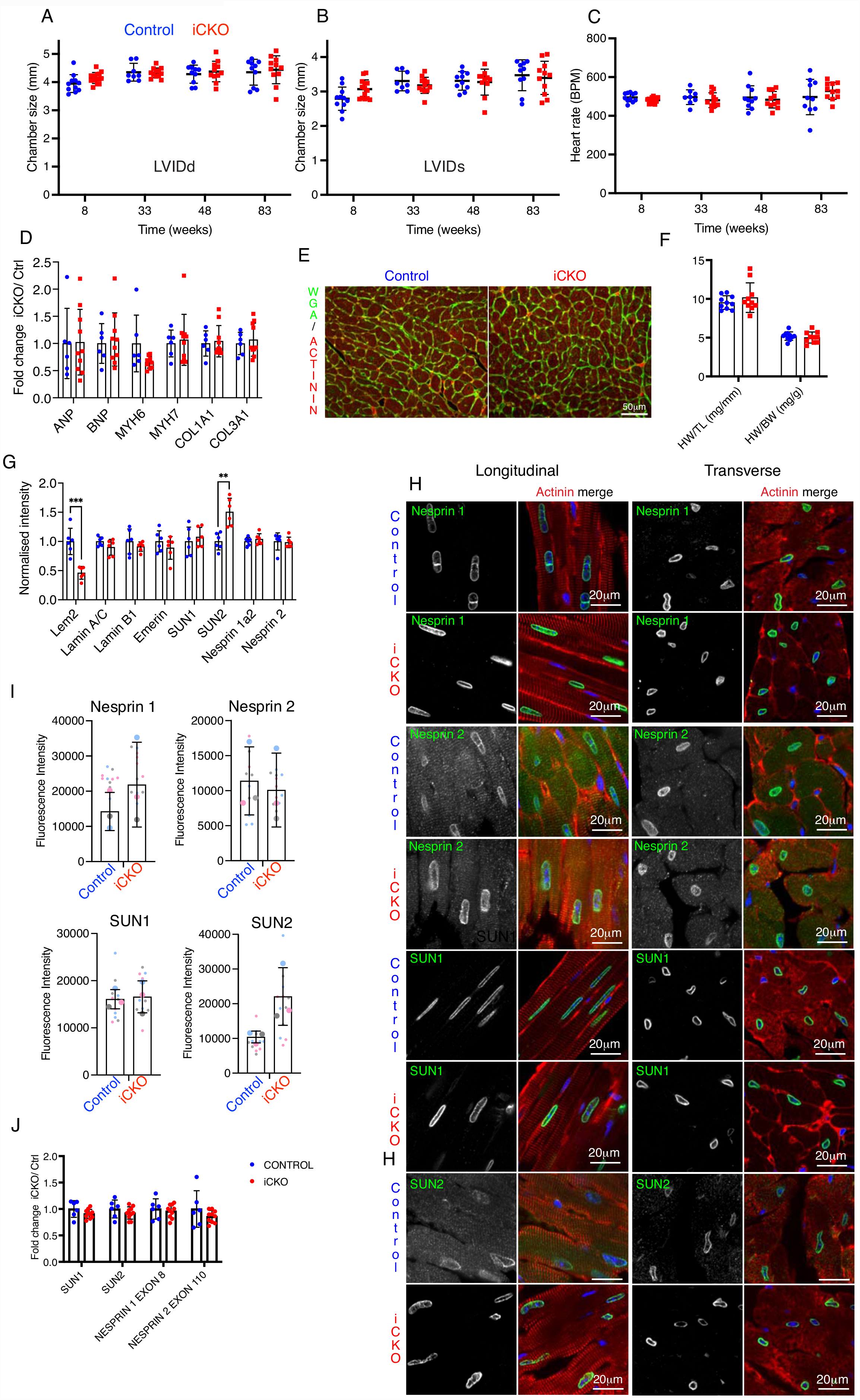
Lem2 ablation in adult cardiomyocytes does not affect heart function or lead to adverse cardiac remodelling. Echocardiography was performed on Lem2f/f (control) and Lem2f/f;Cre/+ (cardiomyocyte-specific tamoxifen-inducible Lem2 knockout, iCKO) at 8 weeks of age, one week prior to tamoxifen injection, and subsequently followed up at 33, 48 and 83 weeks of age. Measured parameters were as follows: **(A**,**B)** Left Ventricle Internal Diameter during diastole (LVIDd) and systole (LVIDs); and **(C)** heart rate. Note that as expected, several effects of ageing were evident, including increased LVIDd and LVIDs (A,B), but no differences were observed in any of the parameters between control and iCKO animals, at any time point. **(D)** Quantitative real-time PCR was performed using primers designed to amplify fetal gene programme markers (ANP, BNP, Myh6, Myh7) and pro-fibrotic genes (Collagen 1a1 and 3a1) on RNA extracted from 83 week old mouse hearts. Fold changes were calculated and compared between iCKO and Control mice. No significant differences were observed between the genotypes indicating there is no adverse remodelling nor fibrotic responses in the iCKO mice. **(E)** Heart sections from 83 week-old mouse hearts were stained with fluorescently-labelled wheat germ agglutinin (WGA, green)/α-actinin (red) to delineate cell boundaries and cardiomyocytes, respectively (lower panels). **(F)** Heart weight: tibia length (HW/ TL) and heart weight: body weight (HW/ BW) ratios in 83 week-old control and iCKO mice, showing no differences. **(G)** Quantification of Western blots from Figure 5G. **(H**,**I)** Immunofluorescence on sections of 83-week old hearts, stained with antibodies raised against LINC complex components Nesprin 1, Nesprin 2, SUN1, SUN2 (green), in both the longitudinal (left panels) and transverse planes (right panels) of cardiomyocytes, quantified in (I). Large data points in (I) represent means from individual mice, small points represent technical replicates within each mouse. **(J)** qRT-PCR of LINC complex genes Nesprin 1 (exon 8) and Nesprin 2 (exon 110), SUN1, SUN2. **p<0.01, ***p<0.001, two-tailed t-test; n=10-14 mice per genotype for echocardiography, n=6-10 for qRT-PCR, n=6 for western blots, n=3 for immunohistochemistry. Graphs show mean +/-standard deviation.

**Supplementary Figure 5:**
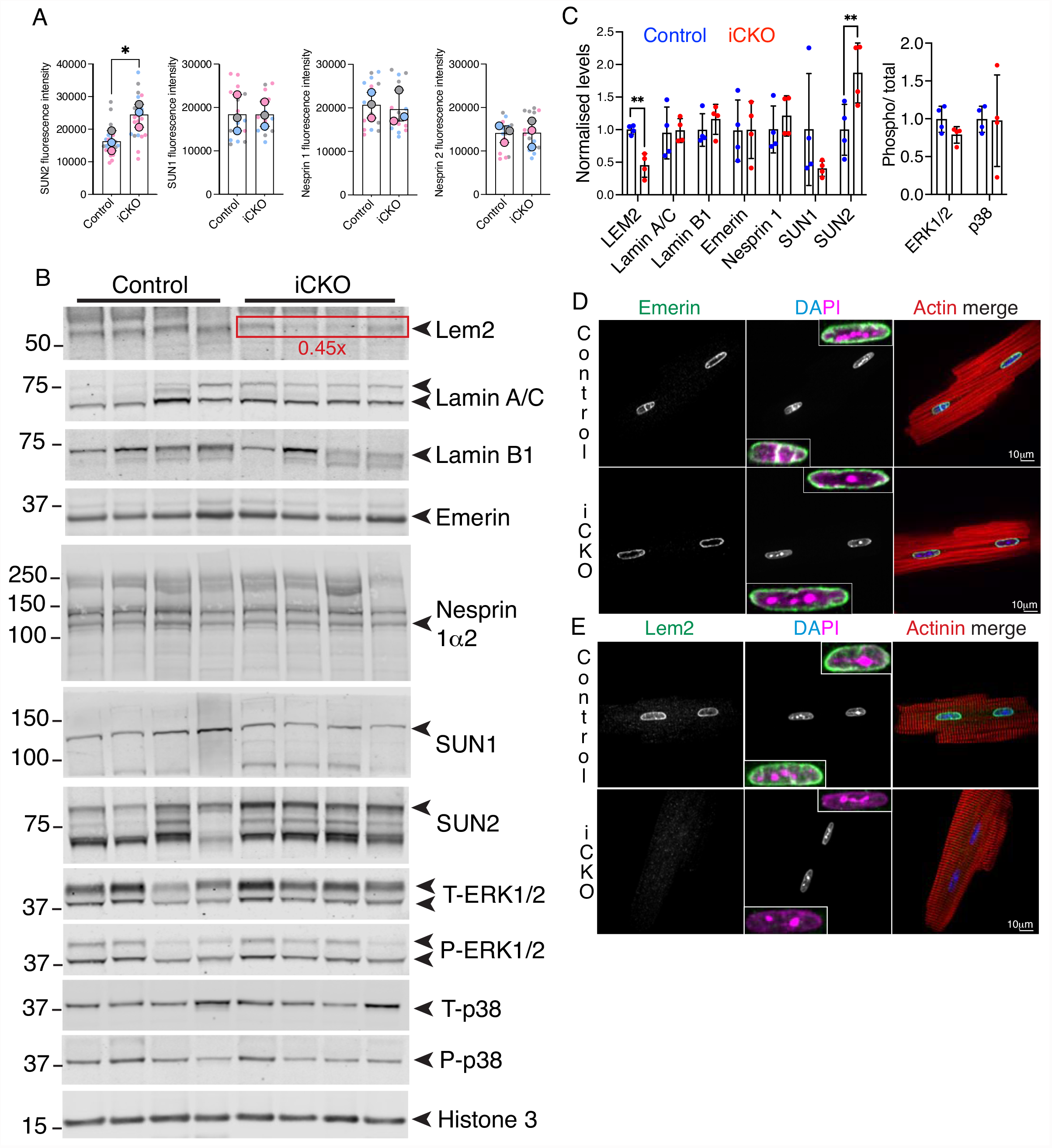
Levels of nuclear lamins, nuclear envelope, and LINC complex proteins in adult cardiomyocytes isolated from Lem2 iCKO mice. **(A)** Quantification of LINC complex fluorescence intensity levels from Figure 5A. **(B)** Western blots were performed on protein extracted from isolated adult cardiomyocytes from 8-10 week-old mice using antibodies raised against Lem2, Lamins A/C, B1, Emerin, SUN1, 2, Nesprin-1α2, phospho-ERK1/2, total ERK1/2, phospho-p38, total p38 and Histone H3 (loading control). **(C)** Quantification of (B). Note the decreased levels of Lem2 in Lem2 iCKO compared to control. n=4 mice per genotype. **(D**,**E)** Adult cardiomyocytes isolated from 8–10-week-old control and Lem2 iCKO mice were stained using antibodies raised against Emerin (D) and Lem2 (E) (green), sarcomeric α-actinin or phalloidin (red) and DNA (blue/ magenta). (A) Large data points represent means from individual mice, small points represent individual nuclear measurements within each mouse. n=3 mice/genotype for immunohistochemistry, n=4 for Western blot. Graphs show mean +/-standard deviation.

## Supplementary Movies

**Supplementary Movie 1: E16.5 HREM ventral-dorsal erosion**. Control (left) and cardiomyocyte-specific Lem2 knockout (right) hearts were imaged using HREM and eroded from ventral to dorsal. Note the thinner walls observed in the cKO hearts, but apparently normal formation of the atrial and ventricular septa, and atrio-ventricular connections. Movies are representative of n=5-7 hearts.

**Supplementary Movie 2: E16.5 HREM base-apex erosion**. Control (left) and cardiomyocyte-specific Lem2 knockout (right) hearts were imaged using HREM and eroded from base to apex. Movies are representative of n=5-7 hearts.

**Supplementary Movie 3: E14.5 HREM ventral-dorsal erosion**. Control (left) and cardiomyocyte-specific Lem2 knockout (right) hearts were imaged using HREM and eroded from in ventral to dorsal axis. Note the slightly thinner walls observed in the cKO hearts, but apparently normal formation of the atrial and ventricular septa, and atrio-ventricular connections. Movies are representative of n=3-5 hearts.

**Supplementary Movie 4: Live cell imaging of cardiomyocytes expressing photoactivatable Lem2**. Neonatal cardiomyocytes transiently expressing Lem2-mEOS2 were photoactivated and imaged live. Note the highly dynamic structure of the cardiomyocyte nuclear envelope.

**Supplementary Movie 5**

Z stacks of nuclei from control adult cardiomyocytes, labelled with antibodies to Lamin A/C, sarcomeric alpha actinin (magenta), and DAPI (magenta) and imaged using instant structured illumination microscopy. Sequential erosion through the nuclei reveals numerous invaginations in control nuclei.

**Supplementary Movie 6**

Z stacks of nuclei from iCKO adult cardiomyocytes, labelled with antibodies to Lamin A/C, sarcomeric alpha actinin (magenta), and DAPI (magenta) and imaged using instant structured illumination microscopy. Sequential erosion through the nuclei reveals numerous invaginations in iCKO nuclei.

**Supplementary Movie 7**

Z stacks of nuclei from control adult cardiomyocytes, labelled with antibodies to Lamin B1, sarcomeric alpha actinin (magenta), and DAPI (magenta) and imaged using instant structured illumination microscopy. Sequential erosion through the nuclei reveals numerous invaginations in control nuclei.

**Supplementary Movie 8**

Z stacks of nuclei from iCKO adult cardiomyocytes, labelled with antibodies to Lamin B1, sarcomeric alpha actinin (magenta), and DAPI (magenta) and imaged using instant structured illumination microscopy. Sequential erosion through the nuclei reveals numerous invaginations in iCKO nuclei

## Materials and Methods

### Animal Studies

All procedures were performed in accordance with the Guidance on the Operation of the Animals (Scientific Procedures) Act, 1986 (UK Home Office). Mice carrying a knockout-first *Lemd2*^*tm1a*^ allele were generated by blastocyst injection of targeted ES cells (#EPD0240_4_D10, EUCOMM) using standard techniques (*53, 54*). Germ line transmission of *Lemd2*^*tm1a*^ was confirmed by genotyping PCR analysis. Heterozygous *Lemd2*^*+/tm1a*^ mice were bred to the FLPeR deleter strain (*55*), to remove the FRT-flanked knockout-first cassette, generating the *Lemd2*^*tm1c*^ floxed allele in which exon2 was retained flanked by LoxP sites. For the Lem2 cKO mouse line, Lem2 floxed (Lem2^f/f^) were crossed with *Xenopus laevis* Myosin Light-Chain 2 (XMLC2)-Cre mice (*27*) to generate Lem2^f/+^; XMLC2-Cre. Male Lem2^f/+^; XMLC2-Cre mice were crossed with female Lem2^f/f^ mice to generate Lem2^f/f^; XMLC2-Cre (Lem2 cKO), Lem2^f/+^; XMLC2-Cre and Lem2^f/f^ (controls). For the Lem2 cardiac Troponin T (cTnT)-Cre cKO mouse line, Lem2 floxed (Lem2^f/f^) were crossed with cTnT-Cre (*28*) to generate Lem2^f/+^; cTnT-Cre. Male Lem2^f/+^; cTnT-Cre mice were crossed with female Lem2^f/f^ mice to generate Lem2^f/f^; cTnT-Cre (Lem2 cTnT-Cre cKO), Lem2^f/+^; cTnT-Cre and Lem2^f/f^ (controls). For the iCKO line, Lem2^f/f^ mice were crossed with tamoxifen-inducible cardiac-troponin T-Cre (Tnnt2^MerCreMer/+^) mice (*45*) to generate Lem2^f/+^; Tnnt2^MerCreMer/+^ mice. Male Lem2^f/f^; Tnnt2^MerCreMer/+^ mice were crossed with Lem2^f/f^ females to generate Lem2^f/f^; Tnnt2^MerCreMer/+^ (Lem2 iCKO) and Lem2^f/f^; Tnnt2^+/+^ (control). Tamoxifen dissolved in peanut oil was intraperitoneal injected for 3 consecutive days at a dose of 0.1mg/g body weight, at 6- and 9-weeks of age for isolated cardiomyocyte and echocardiography studies, respectively.

### Echocardiography

Mice were anesthetized with 0.5% isoflurane and underwent echocardiography using VisualSonics, SonoSite FUJIFILM, Vevo 3100 ultrasound system with a linear transducer 32-55MHz as described previously (*56, 57*). Measurements of heart rate (HR), left ventricular internal dimension at end of diastole and systole (LVIDd, LVIDs, respectively), end-diastolic interventricular septal thickness (IVSd) and LV posterior wall thickness (LVPWd) were determined from the LV M-mode tracing. Percentage fractional shortening (%FS) was used as an indicator of systolic cardiac function.

### Cardiomyocyte isolations and analysis

Adult cardiomyocytes were isolated as described previously (*58*). Fetal cardiomyocytes were isolated using HBSS supplemented with Trypsin (0.5 mg/ml) for 2.5 hours with gentle rotation at 4°C. Hearts were then agitated at 150 rpm on an orbital shaker for 10 mins at 37°C. Trypsin was inhibited with an equal volume of cold culture medium (3:1 mixture of DMEM:M199 media with 4.5 g/l glucose, 10% horse serum, 5% fetal calf serum, 10 mM HEPES and penicillin/streptomycin). Hearts were then disaggregated gently with a 1ml pipette, filtered with a 100 μm cell sieve, pelleted at 150 xg for 5 mins, and resuspended in culture medium. Cells were then plated on collagen/laminin-coated glass. For experiments with 6 kPa PDMS hydrogels, PDMS was prepared and spin-coated onto coverslips as previously described (*38*). Fibronectin coating was used for comparison between glass and hydrogels. Adenoviral transduction of cGAS-mScarlet probe was carried out at 4 infectious particles per cell, to identify nuclear ruptures. Cardiomyocytes were treated with 20μM para-nitroblebbistatin, 5μM verapamil or DMSO (vehicle alone) for 4 hours prior to fixation.

For 2D nuclear shape analysis, the DAPI channel was thresholded by pixel intensity in ImageJ/FIJI, to highlight the full area of each nuclei, followed by measurement using inbuilt algorithms. For semi-quantitative analyses, abnormal nuclear shapes were defined as those showing a bleb/herniation or those showing a concavity in the surface. For 3D shape analysis, z-stacks of images were imported into Volocity software (Perkin Elmer), followed by thresholding of the DAPI channel in 3 dimensions, and measurement using inbuilt algorithms.

### Immunofluorescence

Hearts were fixed overnight in 4% PFA and cryoprotected using sucrose gradients followed by embedding in a sucrose/optimal cutting temperature (OCT) mix as previously described (*59*). 10µm-thick sections were cut and stored at −80°C until use. After permeabilization with washing buffer (PBS with 0.2% TX-100) sections were incubated with the indicated antibodies (Supplementary Table 1) overnight in blocking buffer (PBS/2% normal donkey serum/3% BSA) in a humidified chamber at 4°C. Sections were rinsed in wash buffer and incubated at room temperature with the indicated fluorescently-conjugated secondary antibodies and DAPI (1:1000) diluted in blocking buffer for 1h. Slides were rinsed in wash buffer and mounted in mounting medium (Dako). Sections were imaged as described previously (*60, 61*). 200µL 5-Ethynyl-2’-deoxyuridine (EdU, Molecular probes; 3g/L) was injected intraperitoneally in pregnant females 2 hours before embryo isolation.

### Fluorescence Intensity Analysis

For fluorescence intensity analysis of lamina, NE and LINC complex proteins, line profiles/scans were drawn in ImageJ/FIJI to bisect nuclei. Using the relevant fluorescent channel for each stain, peak intensities corresponding to the nuclear periphery were taken, and the cytoplasmic background signal was subtracted.

### HREM analysis

Samples were prepared as described previously (*62*). Brightness/ contrast adjustments were made using Adobe Photoshop and images assembled using Horos software. Wall thickness measurements were taken at 2 points using sections generated *in silico* by eroding from the base of the heart: at the apex when both ventricles become visible and at the mid-point between the apical section and the top of the ventricles (examples shown in Figure 1D and SF2B). The measurements were then made relative to their respective axis section width and length, and mean values reported.

### Western blotting

Ventricles were lysed in 50µl/mg tissue in a Precellys homogenizer and sonicated to shear DNA. Lysates were centrifuged at 13,900g at 4°C for 15 mins, supernatants aspirated and loaded on to 4-12% NuPage Bis/Tris gels for separation before transfer on to a Nitrocellulose membrane (Biorad) at 4°C at a constant current of 350mA in transfer buffer (25 mM Tris, 190 mM Glycine, 20% Methanol, pH 8.3) for 90 mins. After blocking for 1 hour in Tris Buffered Saline (TBS) containing 0.1% Tween 20 and 5% non-fat dry milk, membranes were incubated overnight at 4°C with the indicated primary antibody (Supplementary Table 1) in blocking buffer (TBS/0.1% Tween 20/2% non-fat dry milk). Blots were washed and incubated with fluorescent secondary antibodies for 1h at room temperature and imaged with an infrared imaging system (ODYSSEY CLx; LI-COR Biosciences). Image Studio software (LI-COR Biosciences) was used for quantitative densitometry analysis to evaluate protein expression levels.

### Histology

Hearts were isolated from age and sex matched littermates at 83 weeks of age, and washed in PBS before being fixed overnight in 10% Formalin. Hearts were subsequently dehydrated in 70% Ethanol, embedded in paraffin and cut into 7 μm sections. Sections were stained using H&E or picrosirius red, mounted and imaged on an All-in-one fluorescence microscope (BZ-X700, Keyence).

### Real-time PCR

Total RNA was extracted from mouse hearts using ReliaPrep™ RNA Tissue miniprep system (Promega). cDNA was synthesized using M-MLV Reverse Transcriptase (Promega). Primers for RT-PCR are listed in Supplementary Table 2. RT-PCR reactions were performed using SYBR-Green (PCR biosystems) in a StepOnePlus Real-time PCR System (Applied Biosystems).

### RNA sequencing

RNA was isolated and its quality assessed using a Bioanalyser (2100 Instrument, Agilent). High quality RNA (RIN score>8) was used for library preparation. 1mg of RNA was used to generate sequencing libraries using NEBNext Ultra II Directional RNA library Prep Kit for Illumina (NEB) according manufacturer’s recommendation. Purification steps were perfomed using the “ProNex Size-Selective Purification System (fragment cut-off size 100bp). The index primers employed during the library preparation corresponded to the ‘NEBNext Multiplex Oligos for Illumina (Index Primers Set 1)’. Once the libraries were generated, their quality was evaluated on the Bioanalyzer. Those libraries whose electropherograms showed a narrow distribution with a peak size of approximately 300 bp, were sent for sequencing. Paired-end multiplexed sequencing of libraries to generate reads of 100 bp was performed on a HiSeq 1000 instrument with TruSeq SBS and PE Cluster v3 Kits (Illumina); 70–100 million reads per sample were obtained. A minimum of 4 hearts per genotype were used for sequencing and analysis. Data will be made available in the European Molecular Biology Laboratory–European Bioinformatics Institute (EMBL-EBI) database.

### RNA-Sequencing data analysis

The raw read counts (FastQ files) were trimmed and aligned (STAR) to the UCSC *Mus musculus* reference genome using Partek Flow software. The gene counts obtained were normalised (GSA) and differential expression analysis was performed applying the DESeq2 method. For further analyses, *p*-values were corrected for multiplicity and the threshold used was false discovery rate (FDR) ≤ 0.1.

### Functional annotation

Gene set enrichment analysis (GSEA) was performed using fgsea package (*63*), version 1.10.1 and molecular signatures database (MSigDB) version 6.2 (*64*) on all genes that were expressed above background levels (readcounts > 600) with settings ‘minSize=15’, ‘maxSize=500’, and ‘nperm=10000’. *P* < 0.05 was considered significant.

### Molecular biology

pTRIP-CMV-GFP-FLAG-cGAS E225A-D227A was a gift from Nicolas Manel (Addgene plasmid # 86674; http://n2t.net/addgene:86674; RRID:Addgene_86674), and Lem2 were subcloned into mScarlet-1 and mEOS-C1, respectively, using XhoI restriction enzyme, then into pShuttle-CMV with EcoRV using the In-Fusion HD Cloning Plus System kit (Takara Bio USA). These were linearized with Pme1 and cotransformed with pAdEasy-1 into BJ5183 cells. Restriction digest and DNA sequencing identified correct clones, which were transformed into JM109 cells. Purified recombinant plasmid DNA was digested with PacI and transfected into HEK293 cells with Lipofectamine (Invitrogen) according to manufacturer’s instructions. Transfected cells were harvested and freeze-thawed, centrifuged at 4000rpm for 10mins and supernatant collected, which was used for subsequent rounds of infection and amplification.

### Adenovirus purification and quantification

Adenovirus purification was performed according to manufacturer’s instructions (Adeno-X Maxi Purification kit). Briefly, HEK293 cells were seeded in 12 wells of a 24-well plate at a density of 2.5×10^5^ cells/mL (1mL per well). The next day, 10-fold serial dilutions (from 10^−2^ to 10^−7^) of the purified viral samples were prepared using DMEM (10%FBS, 2% GPS) and 50 μL of every dilution was added dropwise to each well. Infected cells were incubated at 37°C and 5% CO2 and after 48h, titration procedure was carried out by counting the number of fluorescent cells in 5 fields per well. The infectious units (IFUs) per mL were calculated using the following formula: [(average number of infected cells/field) x (total fields/well)] / [(amount of viral dilution in mL) x (dilution factor)].

### Statistics

Data are presented as mean ± standard deviation. We used 2-tailed Student’s t test or 2-way ANOVA, with Sadik’s post-hoc test for comparisons among groups as indicated. Analysis was performed using Microsoft Excel or Graphpad Prism software. P values of less than 0.05 were considered significant (*, P<0.05; **, P<0.01; ***, P<0.001, ****P<0.0001).

